# Generation of hepatocyte-like cells from sheep embryonic stem cells

**DOI:** 10.64898/2026.10.23.761310

**Authors:** Fatma O. Kok, Alvile Kasarinaite, Deepika Rajesh, Patrick J. Mee, David C. Hay

## Abstract

Development of in-vitro liver models in livestock species presents new opportunities for studying animal physiology and expanding biotechnological applications. Here, we describe a serum-free differentiation strategy to generate hepatocyte-like cells (HLCs) from sheep embryonic stem cells (ESCs). The HLCs displayed hepatocyte morphology and key hepatic functions such as albumin secretion, inducible P450 activity, lipid uptake and storage. Transcriptomic analysis further confirmed progressive hepatic specification and revealed both conserved and species-specific gene expression patterns. Overall, this study establishes a robust system for generating sheep HLCs and offers a platform for studying liver biology in livestock and exploring further biotechnological applications, including cellular agriculture.

## Introduction

Global demand for meat continues to rise alongside population growth and increasing per capita protein consumption. At the same time, conventional livestock production is associated with substantial environmental burden, zoonotic disease risks, ethical concerns and socioeconomic issues (Gerber 2013; Scollan et al. 2010). Therefore, there is an urgent need for methods for protein production that are more sustainable and animal welfare conscious to ensure global food security. The emerging field of cellular agriculture aims to produce animal derived food products using fewer animals than the current livestock industry by cultivating edible tissues directly from cells (Barzee et al 2022; Bhat et al 2014). While current research and commercialization efforts are mostly focused on production of cultured meat and fat, other tissues and organ systems are mostly overlooked (Nie et al. 2025; Goswami et al. 2024; Post 2012; Martins et al. 2024; Mathieu et al. 2025; Hatano et al. 2026). Due to its high nutritional value, distinctive flavour profile and cultural significance in many cuisines, cultivated liver tissue is a viable option for supplementation of different food products, including, pâtè, haggis, and foie gras.

A major bottleneck in development of cell-cultivated products is the limited performance of animal cells in-vitro. Most of the cell types derived from livestock in their native state have severe limitations with respect to in vitro culture, presenting major challenges for the scalability and cost-effectiveness of cultivated meat bioprocesses. A good example of this are satellite cells that are traditionally used to generate cultivated meat. They display limited cellular lifespans and slow proliferation rates in-vitro (Post 2012; Thorrez and Vandenburgh 2019). Therefore, other sources of somatic cells have been explored. This includes pluripotent stem cells (PSCs) which display key attributes, including indefinite proliferation potential and the plasticity to differentiate into any tissue type. Due to these properties PSCs have emerged as a viable option for large scale production of cultivated meat without the need for repeated donor organ harvesting and extensive processing (Zhu et al. 2023).

Here, we describe a robust and efficient generation of hepatocyte-like cells (HLCs) from sheep ESCs using a serum-free system. To our knowledge, this is the first time that these cells are shown to be able to differentiate into hepatic lineage. Sheep HLCs exhibit typical hepatocyte morphology, express lineage specific markers and have characteristics of mature hepatocytes such as albumin secretion, lipid uptake and accumulation, and inducible cytochrome P450 activity in-vitro. This differentiation protocol provides a powerful alternative to conventional livestock production of liver tissue and offers promising potential for application in large scale cultured organ meat production. The establishment of ovine liver models will also improve our understanding of basic animal biology, including animal health, complex disease, and aging. All of which have direct impacts on the efficiency of raising livestock, animal welfare, farming economics, and the environment. Additionally, the establishment of a robust differentiation protocols provide future opportunities to develop cost effective scale-up platforms for mass production. This will not only benefit cellular agriculture, and provides prototype templates for the mass production of human tissue for future biomedical application.

## Results

### Differentiation of sheep ESCs into HLCs

Our aim was to develop a liver differentiation process from sheep embryonic stem cells (ESCs). Presently, there key knowledge gaps in the embryonic development of the sheep. Therefore, we modified existing differentiation protocols that had been developed in for humans and other mammals, with a focus on producing the major metabolic cell type of the liver, the hepatocyte (Agarwal et al. 2008; Cheng et al. 2012; Teo et al. 2012; Touboul et al. 2010; Song et al. 2009; Szkolnicka et al. 2014; Medine 2014; Hay et al. 2008; Hannan et al. 2013a, b; Brolen et al. 2010; Argarwal et al. 2008; Yiangou et al. 2018; Pauklin and Vallier 2013). To mimic mammalian development, the differentiation of process was performed in stages, directing ESCs from pluripotency via hepatic progenitor to the hepatocyte lineage. Specifically, the three-stage process involved the induction of definitive endoderm (DE), then hepatic progenitor cells (HPCs), and followed by hepatocyte-like cell (HLC) specification.

Although nodal/Activin signalling is known to have central role in vertebrate DE formation, other signalling pathways, such as bone morphogenetic protein 4 (BMP4), are also implicated (Teo et al. 2012; Stainier 2002). Critical to organogenesis, is the correct spatially and temporally control of morphogen exposure (Inge et al. 2025). Therefore, we optimized the dose and duration of Activin A or a combination of Activin A and BMP4 to promote DE differentiation from sheep ESCs. Unlike typical human PSC differentiation protocols where cells should reach 40-50% confluency before initiation of differentiation, sheep ESC differentiation was initiated 24 hours post seeding, when the cells were 10-15% confluent (Fig1A). During DE differentiation, morphology of the sheep ESCs changed dramatically from typical colony morphology to bright, densely packed clusters (Fig1A).

**Fig 1:**
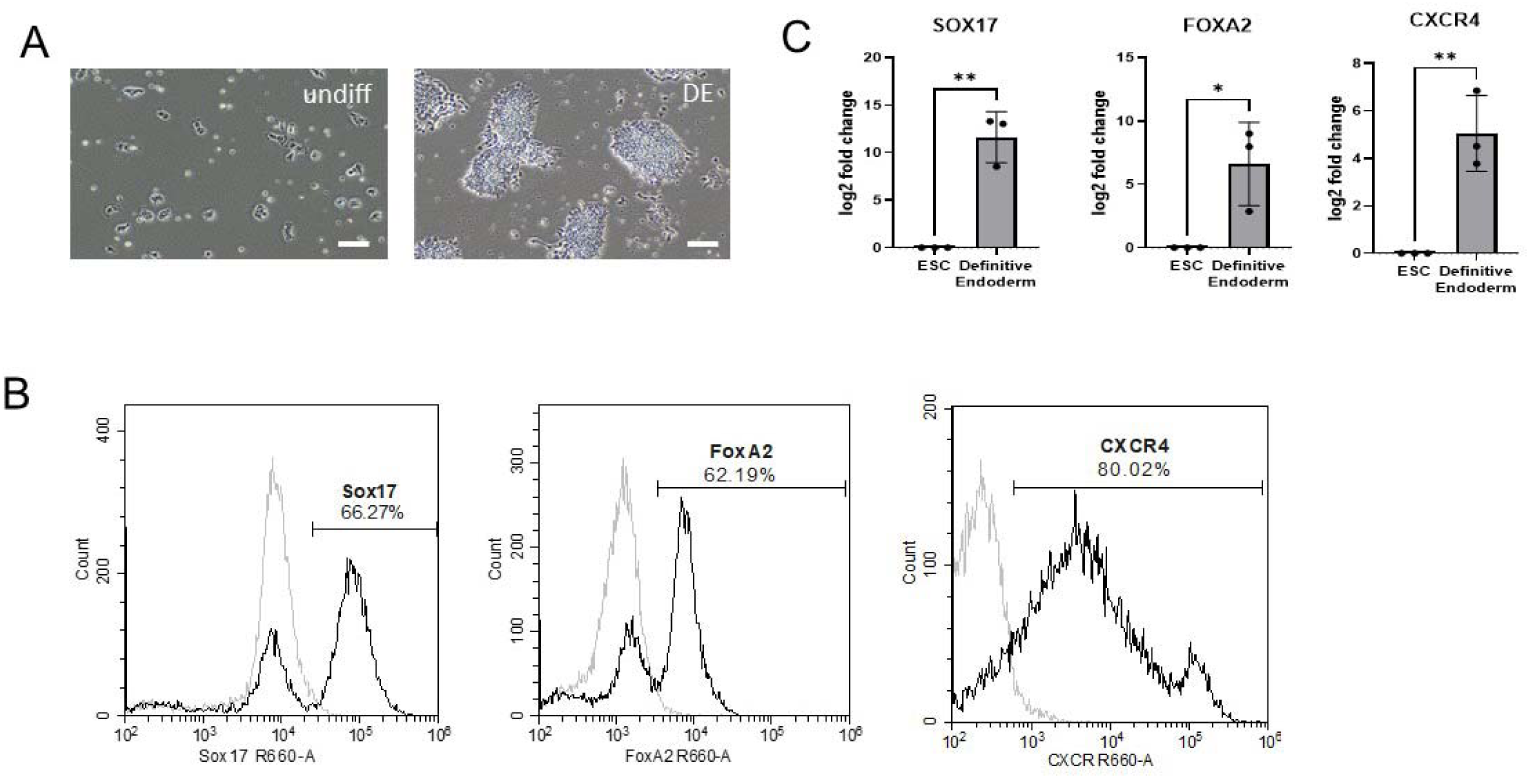
Ovine pluripotent stem cell differentiation to definitive endoderm (DE). A) Representative images showing cell morphology at the start of differentiation (undiff) and at the definitive endoderm stage B) Flow cytometry analysis of Sox17, FoxA2 and CXCR4 expression in DE cells. Gray histogram represents isotype control, black histogram represents indicated antibody staining. C) qPCR analysis of SOX17, FOXA2 and CXCR4 marker expression in DE compared to undifferentiated controls (n=3, p<0.05).

Morphological changes were paralleled by increased expression of developmentally relevant markers. Flow cytometry analysis of differentiated DE cells showed that differentiating cells expressed Sox17 (∼66%), FoxA2 (∼62%), and CXCR4 (∼80%) (Fig1B). This was paralleled by a dramatic decrease in Oct4 expression, from 90% cells at pluripotent state to 33% at DE stage (Suppl Fig 1). DE transcription factor and cell surface protein expression was further supported by qPCR analysis demonstrating the significant upregulation of SOX17, FOXA2, and CXCR4 during DE specification (p<0.05, n=3) (Fig1C).

Following efficient levels of DE specification, we proceeded with HPC differentiation. Dimethylsulfoxide (DMSO) is known to be an important inducer of hepatic specification in human differentiation models (Hay et al. 2008; Siller et al. 2015; Sullivan et al. 2010; Du et al. 2018). Therefore, we used this approach to differentiate sheep DE toward the hepatic lineage. During early stages, tightly packed DE clusters loosened up and became flatter (Fig 2A). At later stages, a wave of proliferation was observed with high number of cells exhibited typical hepatic progenitor like (HPC) morphology (Fig2A). DE transition into the hepatic lineage was followed by flow cytometry and qPCR. Differentiating HPC populations were homogenous with >95% of cells expressing cytokeratins 8/18/19 (ctKRT), and >80% of the cells expressing Hnf4A and FoxA2 (Fig2B). These data were supported by gene expression analysis with increased expression of hepatic genes, HNF4A, albumin (ALB) and alpha fetoprotein (AFP) when compared to undifferentiated ESCs (p<0.05, n=3) (Fig2C).

**Fig 2:**
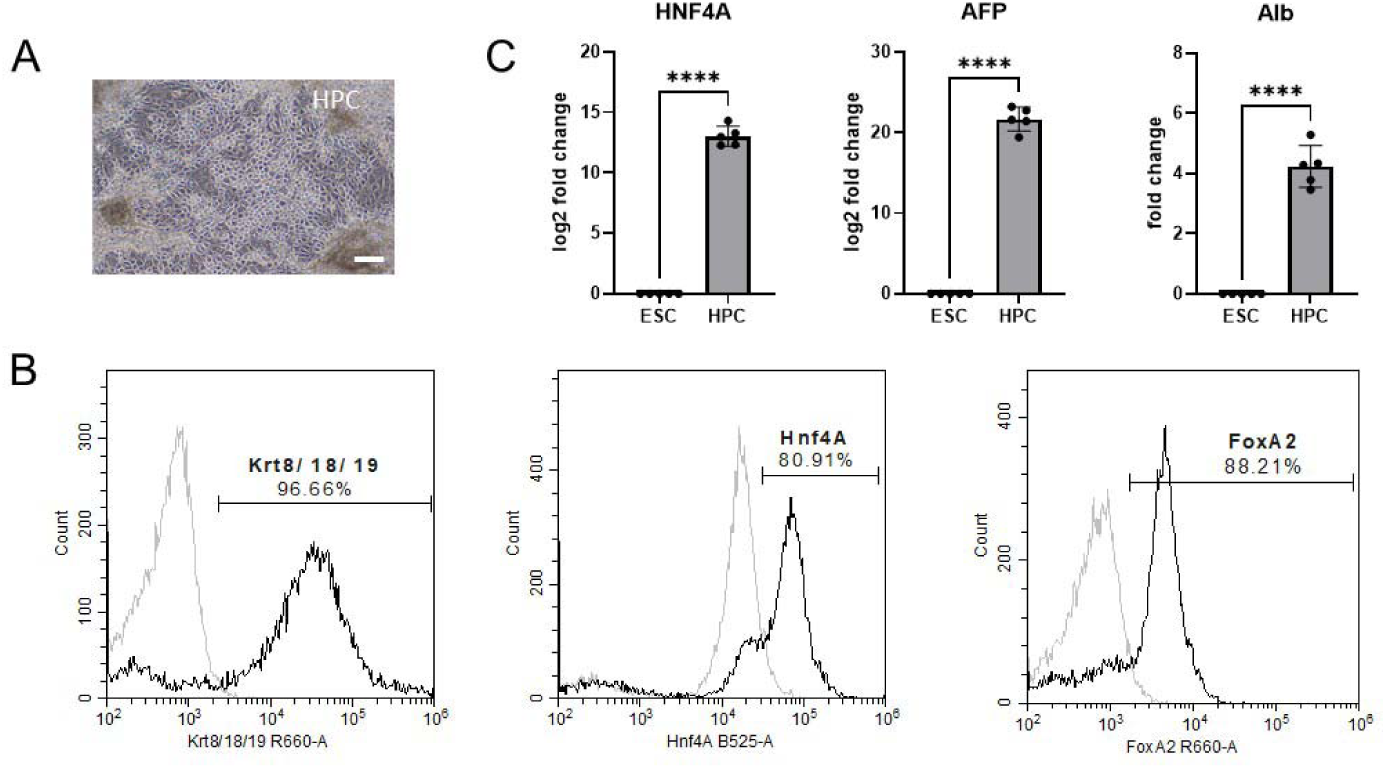
Hepatocyte-progenitor cell (HPC) specification from pluripotent stem cells. A) Representative image showing cell morphology of HPCs. B) Flow cytometry analysis of Hnf4A, FoxA2 and Krt8/18/19 expression in HPCs. Gray histogram represents isotype control, black histogram represents indicated antibody staining. C) qPCR analysis of HNF4A. AFP. ALB marker expression in HPCs compared to undifferentiated controls (n=3, p<0.05).

Given the previous cell transitions were efficient, we proceeded with the maturation of differentiated HPCs into HLCs previously reported media containing hepatocyte growth factor (HGF) and oncostatin M (OSM) (Szkolnicka et al. 2014, 2026; Wang et al. 2017). During maturation, differentiation cells became larger and polygonal with increased cytoplasmic-to-nuclear ratio compared to HPCs (Fig3A). During this process, HLCs retained their AFP expression, in concert with and ALB, indicating further HLC maturation maybe required in future experiments (Rashidi et al. 2018)(Fig3B).

**Fig 3:**
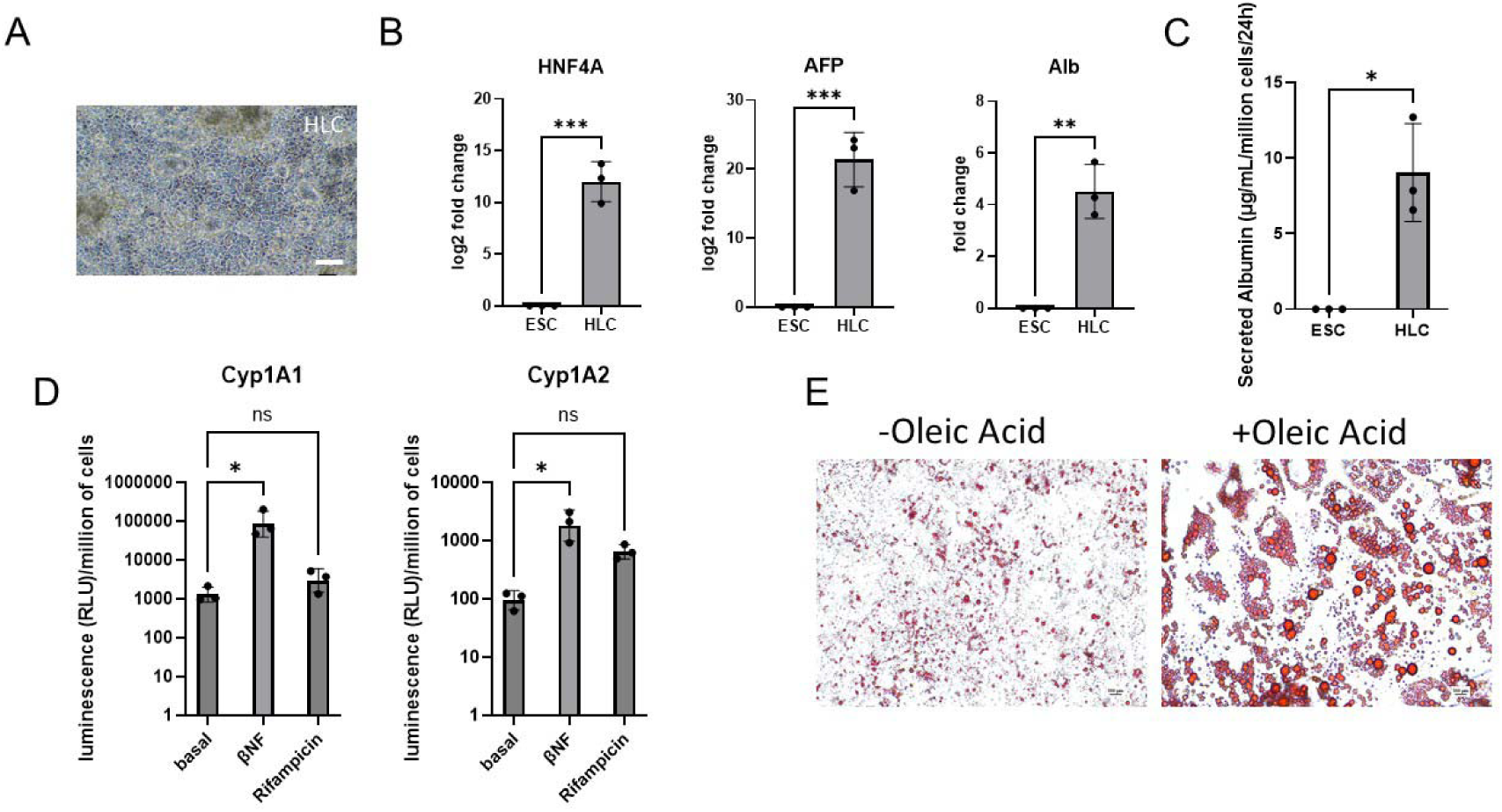
Hepatocyte-like cell (HLC) specification and differentiation. A) Representative image showing cell morphology of HLCs. B) qPCR analysis of HNF4A, AFP, ALB marker expression in HLCs compared to undifferentiated controls (n=3, p<0.05). C) Quantification of albumin secretion in HLCs by ELISA (n=3, p<0.5) D) Basal and induced CYP1A1 and CYP1A2 activity in HLCs following treatment with rifampicin and β-naphthoflavone E) Lipid accumulation in HLCs with and without oleic acid loading and assessed by Oil Red O staining

### Functional characterization of HLCs

To further characterise HLC function and maturity, we performed a range of assays. One key function of hepatocytes, is their ability to produce and secrete serum proteins, such as albumin. To detect albumin secretion, we collected culture medium supernatant samples and measured albumin levels by ELISA. While culture medium collected from undifferentiated ESCs had minimal levels of secreted albumin, sheep HLCs secreted 9.03±1.87 ug/mLalbumin in 24 hours (n=3, p<0.05) (Fig3C). Unfortunately, it was not possible to test for AFP secretion as commercially available kits did not cross react with the ovine protein. Therefore, to test for maturity, we employed another strategy which focussed on drug metabolism by the cytochrome P450 enzyme family (CYP450). We assessed the expression of functional and induction of two developmentally relevant CYP450s. Cyp1A1 which is considered to be active in the fetal liver, but drug inducible in the adult. We also assessed the function and induction Cyp1A2 which is expressed in the neonate and adult. We observed high basal activity of Cyp1A1 and Cyp1A2 indicating a mixture of fetal and adult characteristics in HLC monolayers (Fig3D). Notably, both CYP450 isoforms were drug inducible with β-Naphthoflavone (β-NF) but not rifampicin, demonstrating enzymatic specificity, and mature hepatocyte traits. (Fig3D). Finally, we tested sheep HLC ability to uptake lipids using Oil Red O staining. Lipid storage it is a feature of both fetal and adult liver, with the former associated with developmental energy requirements and the latter generally associated with pathogenesis in adults. The number and size of the lipid droplets were significantly increased following supplementation of maturation medium with Oleic acid, indicating functional uptake and storage of exogenous fatty acids (Fig3E).

### Transcriptomic analysis of sheep hepatic differentiation

To characterize the global transcriptional changes occurring during sheep hepatic differentiation and to evaluate lineage progression and hepatic specification of sheep ESCs in detail, RNA sequencing (RNA-seq) was performed (Fig4). Principal component analysis (PCA) of the RNA-seq data showed clear stage specific clustering with ESCs, DEs, HPLs and HLCs each forming distinct groups. Closer proximity between HPCs and HLCs relative to earlier stages suggests gradual lineage commitment and establishment of hepatic identity (Fig4A). Expression analysis of pluripotency markers revealed a sharp decline at DE stage with complete loss of expression in HPCs and HLCs, confirming progressive loss of stem cell identity during differentiation (Fig 4B).

**Fig 4:**
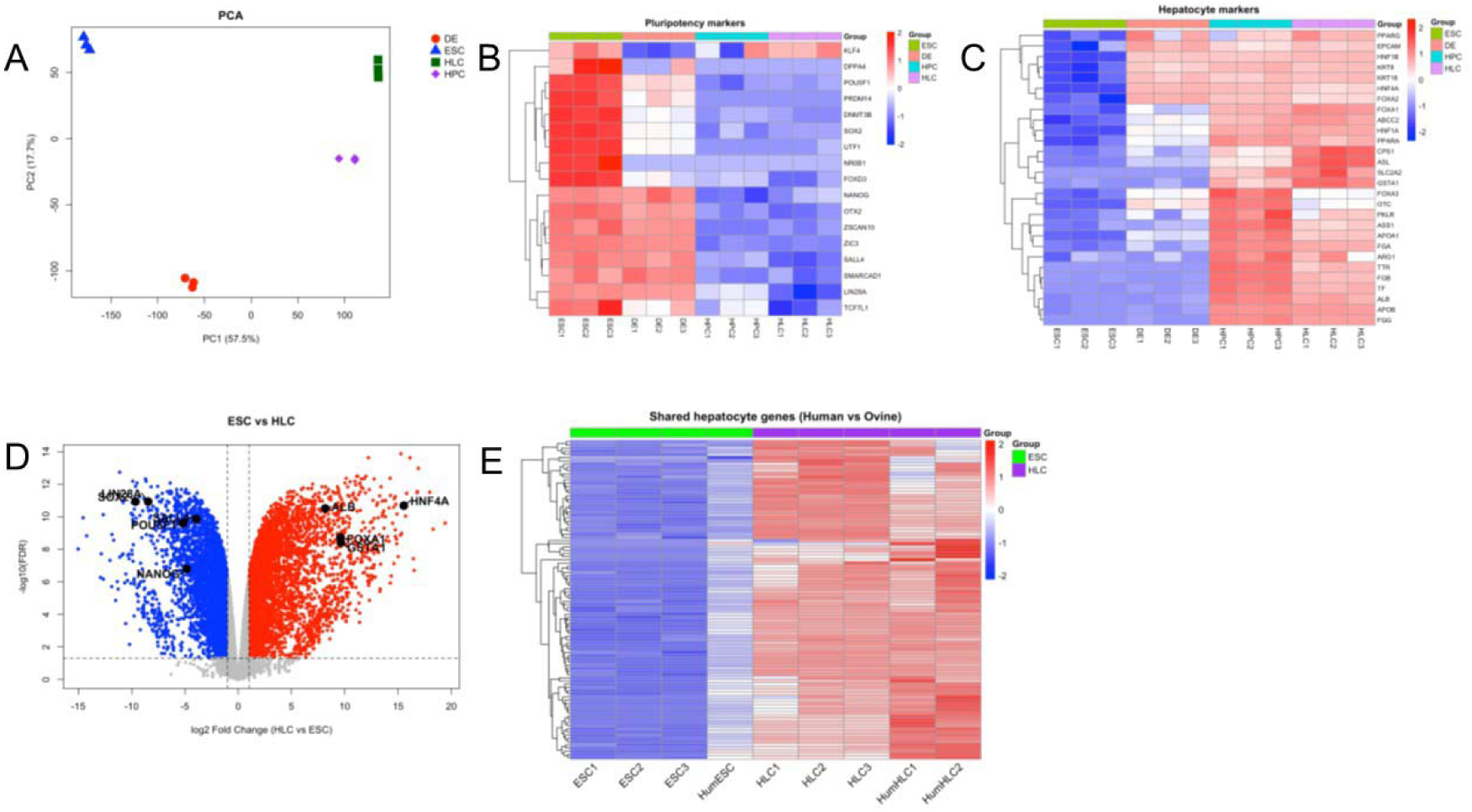
Whole transcriptome analysis from pluripotency to HLC differentiation in sheep. A) Principal component analysis (PCA) of RNA-seq data at different stages of differentiation B) Heatmap of pluripotency marker expression across different stages of differentiation C) Heatmap of hepatocyte marker expression across different stages of differentiation D) Volcano plot showing differential gene expression in HLCs compared to undifferentiated ESCs E) Heatmap comparing gene expression profiles of human and sheep ESCs and HLCs.

Analysis of hepatic lineage associated genes revealed a stepwise activation of hepatic transcriptional program during differentiation. At DE stage, cells began expressing subset of early pioneer transcription factors including HNF4A and FOXA2, indicating early commitment toward hepatic fate. However, expression of late-stage hepatic markers remained largely absent at this stage, indicating that full hepatic specification was not achieved at this point. In contrast, HPCs exhibited robust induction of both early and late-stage hepatic genes, which was maintained or further induced in mature HLCs. Together, these data support the stepwise establishment of hepatic identity through sequential activation of lineage specific transcriptional networks (Fig 4C). Consistent with stage-specific marker analysis, volcano plot comparison between ESCs and HLCs demonstrated significant upregulation of markers specific to hepatic identity, such as HNF4A and ALB while repression of pluripotency associated genes. This highlighted the extensive transcriptional reprogramming that occurred during hepatocellular specification (Fig 4D).

Comparative transcriptomic analysis between sheep and human cells (Ma et al. 2022) demonstrated a core set of genes with highly similar expression profiles, indicating strong conservation of key transcriptional programs underlying pluripotency and hepatic identity. We did, however, identify a subset of genes that exhibited species-specific expression patterns, highlighting divergence in other regulatory programs between two species, despite the overall conservation of lineage-associated gene signatures (Fig 4E, Suppl Fig2). Elevated expression of FGG, FGA, PLG, KNG1, HRG and F5 was observed in ovine HLCs relative to human HLCs. Higher expression levels were also observed for APOB, APOA4, APOH and A1CF. Similarly, the expression of the hepatic identity and secretory genes HNF4A, TTR, SERPINA1 and AHSG was greater in ovine HLCs (Fig 4E, Suppl Fig2)

## Discussion

In this study, we describe a stepwise differentiation strategy for generating HLCs from sheep ESCs and characterize their progression at both functional and transcriptional levels. While hepatic differentiation from human pluripotent stem cells have been extensively studied, little is known about how these developmental transitions are controlled in livestock species. Therefore, developing livestock differentiation models is important not only for understanding species-specific developmental biology, but also for expanding the experimental toolkit available for agricultural biotechnology and cellular agriculture applications. Our findings highlight both conserved and species-specific features of hepatocyte differentiation in vitro. Although many aspects of hepatic lineage progression appeared comparable to those described in well-established human pluripotent stem cell differentiation systems, differentiated cells displayed some functional and transcriptional characteristics that are likely intrinsic to species-specific differences in liver biology. Increased expression of FGG, FGA, PLG, KNG1, HRG and F5 was observed in ovine HLCs and is associated with coagulation and fibrinolytic pathways. Additionally, greater expression of APOB, APOA4, APOH and A1CF was observed, pointing to differences in lipoprotein assembly, lipid transport and lipid metabolism. Similarly, the expression of the hepatic identity and secretory genes HNF4A, TTR, SERPINA1 and AHSG were greater in ovine HLCs. These differences indicate that although the overall hepatic differentiation trajectory was highly conserved between species, species-specific transcriptional programmes remained evident.

In addition to marker expression, the HLCs displayed core functions associated with liver physiology such as albumin secretion, inducible CYP P450 activity, and lipid uptake and storage ability. This is particularly relevant given the hepatic differentiation is often assessed predominantly through marker expression which can overestimate the functional maturity. The combination of transcriptional and functional validation therefore provides stronger evidence that our differentiation protocol produces biologically relevant HLCs, rather than transcriptionally partial intermediates.

To further characterize the differentiation process, we performed whole transcriptome analysis at different differentiation stages. The data revealed clear separation between early hepatic priming and broader hepatic gene activation. The induction of pioneer transcription factors such as HNF4A and FOXA2 at the definitive endoderm stage suggest that hepatic identity is established relatively early, before the wider hepatic programme is activated. This staged progression is consistent with developmental models in which chromatin accessibility and lineage competency control the degree of transcriptional commitment.

Beyond validating the differentiation trajectory, the RNA-seq data can provide a useful framework for refining culture conditions to better align with the transcriptional state of the cells and can be used as a platform for differentiation optimization. Moreover, the cross-species comparison can provide additional insight for species-specific refinement. Although many core pluripotency and hepatic genes showed similar expression patterns between sheep and human cells, the presence of species-specific transcriptional differences suggest that direct transfer of human differentiation protocols may overlook important regulatory distinctions.

Interpretation of our sheep hepatocyte differentiation and maturation process benefited from the increasing availability of species-specific reagents and functional assays. Many commonly used characterization approaches for human PSC derived hepatocytes including antibodies, luminescence based CYP450 assays and ELISAs have been extensively optimized for human systems and provide robust and well-established readouts. When applied in sheep hepatocytes, some of the assays required optimization or were not directly transferable, reflecting expected differences in assay development across species. Collectively, our data results highlight the importance of interpreting hepatocyte differentiation and maturation within a species-specific perspective rather than relying exclusively on human hepatic markers, assays and functional activity.

Beyond its use as a differentiation model, establishing a readily available sheep hepatocyte system opens up broader opportunities in both animal biology and cellular agriculture. For example, the lack of species-appropriate in-vitro hepatocyte models has long been a limitation in veterinary pharmacology and feed additive safety assessment. Therefore, a renewable source of sheep HLCs can help to address this gap, enabling more relevant metabolic and toxicity screening without the ethical and logistical limitations of primary hepatocyte isolation. Moreover, the differentiation strategy developed here provides a foundation for the emerging cultivated meat sector, particularly for production of edible liver tissues with specific nutritional characteristics, without the need of conventional organ harvesting. This approach will also reduce substantial environmental burden, zoonotic disease risks and ethical concerns typically associated with conventional livestock production.

In summary, this work highlights the value of species-specific hepatocyte models, providing a versatile resource which links animal biology and welfare with sustainable food production and the biomanufacturing industry.

## Materials and Methods

### Sheep ESC culture and differentiation

Sheep ESCs (Silva-Almeida et al. 2026) were maintained on laminin-511 (Takara) in N2B27 medium supplemented with 20 ng/ml activin A (R&D Systems), 12.5 ng/ml FGF2 (Peprotech) and 2 µM XAV939 (Merck) (Kinoshita et al. 2021). Cells were passaged using TrypLE (Gibco) when colonies reached 70-80% confluency.

For definitive endoderm induction, 10-15% confluent sheep ESCs were incubated in RPMI 1640 medium (Merck) supplemented with 2% B27 (without Vitamin A) (Gibco), 100ng/mL Activin A (STEMCELL Technologies) and 50ng/mL BMP4 (R&D Systems) for 24 hours. On the following 4 days, medium is supplemented with only Activin A.

For hepatic specification, sheep DE cells were cultured in Knockout-DMEM medium (Gibco) supplemented with 20% knockout serum replacement (Gibco) and 1% DMSO (Merck) for 5 days.

For hepatocyte maturation, sheep hepatoblasts were cultured in HepatoZYME medium (Gibco) supplemented with 10ng/mL Hepatocyte Growth Factor (HGF) (Peprotech), 20ng/mL Oncostatin M (OSM) (Peprotech) and 100nM hydrocortisone 21-hemisuccinate sodium salt (HCC) (Merck) for 6-10 days.

### Quantitative PCR (qPCR) analysis

RNA was extracted from pelleted cells using RNeasy Plus mini kit (Qiagen) according to manufacturer’s instructions. 1ug of RNA was reverse transcribed into cDNA using Affinity Script QPCR cDNA Synthesis Kit (Agilent) according to manufacturer’s instructions using oligodT primers included in the kit. qPCR is performed and analysed using QuantStudio 3 system (Applied Biosystems) using Power SYBR Green PCR master mix (Applied Biosystems). GAPDH was used for internal normalization. Primer sequences are included in Table 1.

**Table 1:**
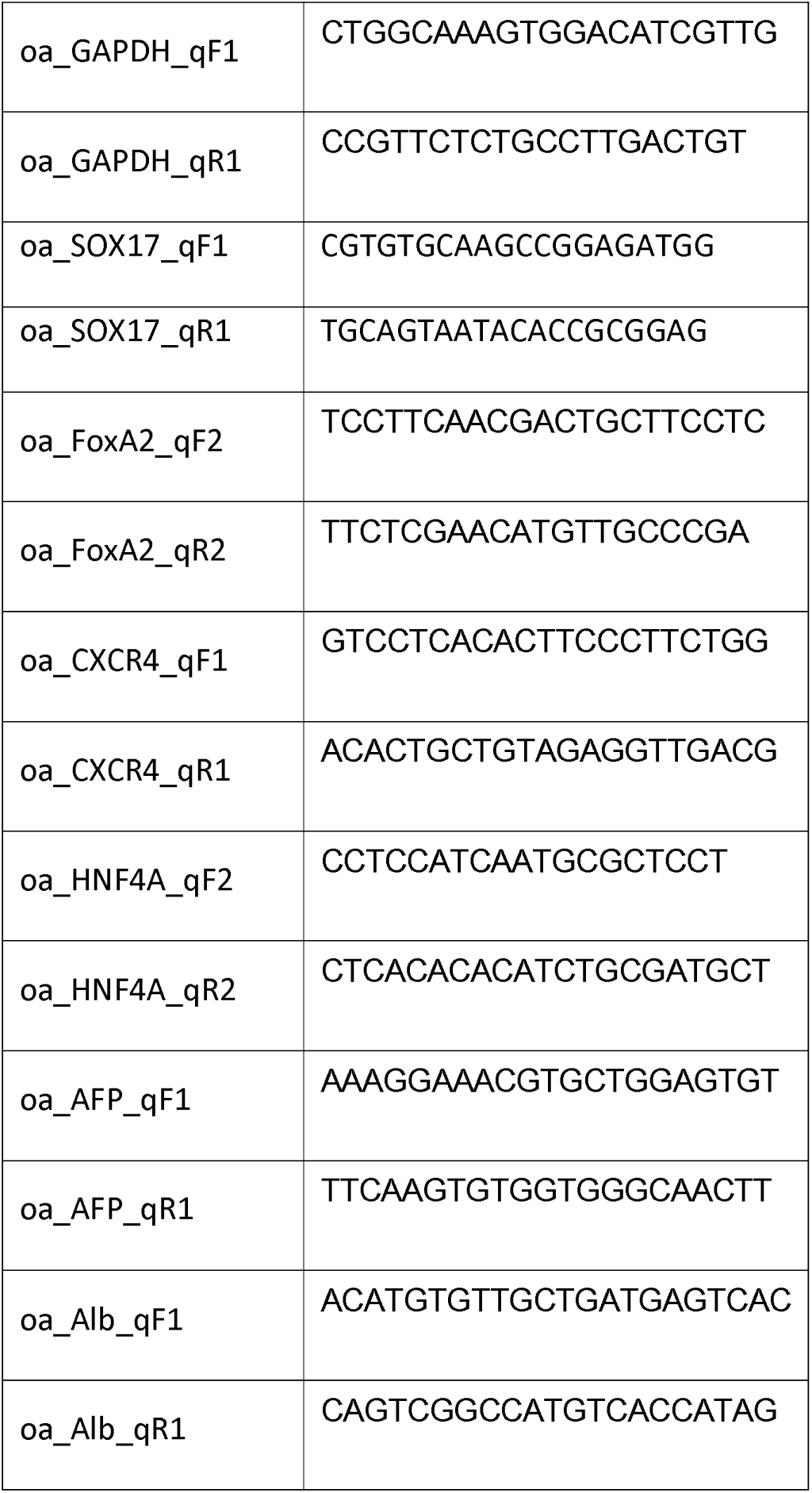
PCR Primers.

### Flow cytometry analysis

Cells were harvested as single cells using TrypLE, resuspended in FACS buffer (dPBS (Gibco) supplemented with 2% FBS (Gibco)) and stained with Live/Dead fixable violet dead cell stain kit (Invitrogen) for 15 mins. For surface marker analysis (CXCR4), cells were stained in FACS buffer for 1 hour immediately after live/dead staining. For internal marker analysis, cells were fixed in 4% PFA (Thermo Scientific) for 15 minutes, washed once in FACS buffer and resuspended in permeabilization buffer (FACS buffer supplemented with 0.1% Triton X-100 (Merck)) for 30 minutes. Cells were then stained using primary antibody in permeabilization buffer for 1 hour in dark. If non-conjugated primary antibody was used, cells were washed three times in permeabilization buffer before staining with conjugated secondary antibodies for 30 minutes in dark. After staining, cells were washed three times in FACS buffer and analysed with CytoFlex (Beckman Coulter) and associated software, CytExpert. The list of antibodies used in this study is included in Table 2.

**Table 2:** Antibodies.

| Antibody | Vendor | Cat number | Final concentration<br>(µg/mL) |
| --- | --- | --- | --- |
| Anti-OCT4 antibody (Rabbit IgG, PE) | Abcam | ab283731 | 0.32 |
| Isotype Control for OCT4 (Rabbit IgG, PE) | Abcam | ab209478 | 0.32 |
| Anti-SOX17 antibody | R&D systems | AF1924 | 2 |
| Isotype Control for SOX17 (Goat IgG) | Invitrogen | 02-6202 | 2 |
| Anti-HNF4A antibody | Santa Cruz | sc-374229 | 2 |
| Isotype Control for HNF4A (Mouse IgG1, kappa) | Abcam | ab170190 | 2 |
| Anti-FOXA2 antibody (human IgG1, APC) | Miltenyi | 130-124-043 | 2 |
| Isotype Control for FOXA2 (human IgG1, APC) | Miltenyi | 130-113-434 | 2 |
| Anti-Cytokeratin 8/18/19 (human IgG1, APC) | Miltenyi | 130-112-347 | 2 |
| Isotype Control for Cytokeratin 8/18/19 (human IgG1, APC) | Miltenyi | 130-113-434 | 2 |
| Anti-CXCR4 (human IgG1, APC) | Miltenyi | 130-120-778 | 2 |
| Isotype Control for CXCR4 (human IgG1, APC) | Miltenyi | 130-113-434 | 2 |
| Secondary Antibody for SOX17 (anti-goat IgG, AF647) | Invitrogen | a32849 | 2 |
| Secondary Antibody for HNF4A (anti-mouse IgG, AF488) | Invitrogen | a21202 | 2 |

### Albumin secretion ELISA assay

Albumin content in the spent media is determined using sheep albumin ELISA kit (Cusabio) according to the manufacturer’s instructions. To account for background signal, absorbance values from blank wells were subtracted from all samples. In addition, values obtained from unspent culture medium were subtracted to correct for baseline signal originating from media components. Secreted Albumin levels were then calculated using a standard curve generated from known albumin standards and normalized as indicated. Undifferentiated sheep ESCs were used as negative control.

### CYP induction and CYP activity assay

Basal and induced CYP450 activities were measured using commercially available cell-based P450-Glo™ assays (Promega) using manufacturer’s instructions. For CYP450 induction, HLCs on day 6 of maturation was incubated in complete maturation media supplemented with either 10µM β-naphthoflavone (MedChemExpress) or 25µM Rifampicin (Sigma Aldrich) for 48 hours. DMSO (Merck) was used as vehicle control. Undifferentiated sheep ESCs were used as negative control.

### Lipid loading and Oil Red O staining

For lipid loading, HLCs on day 6 of maturation was incubated in complete maturation medium supplemented with 500µM Oleic acid (Sigma Aldrich) for 48 hours. For Oil Red O staining, cells were washed with dPBS (Gibco) twice, then fixed with 4% PFA (Thermo Scientific). Cells then washed with 60% isopropanol twice and stained with Oil Red O (Sigma Aldrich) in dark for 15 minutes. Cells then washed with distilled water twice and imaged using Nikon Eclipse Ts2.

### RNA sequencing and bioinformatic analysis

Total RNA was isolated from sheep ESCs, definitive endoderm (DE), hepatic progenitor cells (HPCs) and hepatocyte-like cells (HLCs). RNA was extracted from pelleted cells using RNeasy Plus mini kit (Qiagen) according to manufacturer’s instructions. RNA samples were submitted to Biomarker Technologies (BMKGene, UK) for quality assessment, library preparation and high-throughput sequencing. RNA integrity was assessed prior to library preparation.

RNA sequencing was performed on three biological replicates per differentiation stage (n = 3). RNA sequencing libraries were prepared and sequenced using the Illumina NovaSeq 6000 platform with paired-end sequencing (150bp reads). Adapter sequences and low-quality bases were removed prior to alignment. Processed reads were aligned to the Ovis aries ARS-UI_Ramb_v2.0 reference genome using STAR (Dobin et al. 2013). Gene-level read counts were generated using feature Counts. Downstream analyses were performed in R Studio. Lowly expressed genes were removed using standard expression filtering criteria prior to differential expression analysis. Differential gene expression analysis was performed using the edgeR package, and statistical significance was determined using the Benjamini–Hochberg method to control the false discovery rate (FDR). Genes with an FDR < 0.05 were considered significantly differentially expressed. Principal component analysis (PCA), heatmaps and volcano plots were generated from normalised expression data to assess transcriptional relationships between differentiation stages and to identify lineage-associated transcriptional changes during hepatic specification.

For cross-species comparison, human RNA-seq data were obtained from GEO accession GSE185357 (Ma et al. 2022). Orthologous genes shared between sheep and human datasets were identified and expression changes during hepatic differentiation were compared between species. Shared genes were used to evaluate conserved and species-specific transcriptional programmes associated with hepatocyte differentiation.

### Statistics

Experimental (n=≥3) and control groups (n=≥3) were compared using the Student’s paired t-test. Significance is denoted as *(p<0.05) ** (p<0,01) *** (p<0.001).

## List of Abbreviations

ESC: Embryonic stem cell
PSC: Pluripotent stem cells
DE: Definitive Endoderm
HPC: Hepatic progenitor cells
HLC: Hepatocyte-like cells
BMP4: Bone morphogenic protein
DMSO: Dimethyl sulfoxide
HGF: Hepatocyte growth factor
OSM: Oncostatin M
HCC: Hydrocortisone 21-hemisuccinate sodium salt
CYP450: Cytochrome P450 enzyme family

## Declarations

During the preparation of this work, author(s) used ChatGPT to improve grammar and readability. After using this tool, the author(s) reviewed and edited the content as needed and take full responsibility for the content of the publication.

## Ethics approval and consent to participate

Sheep embryos used in this study were obtained from Abreeds (UK) as surplus frozen embryos generated through routine commercial sheep breeding activities. The embryos were produced under standard agricultural husbandry and breeding practices in the United Kingdom and were not generated as part of regulated animal research requiring a UK Home Office licence under the Animals (Scientific Procedures) Act 1986. No live animal procedures were undertaken by the authors for the purposes of this study.

## Consent for publication

All authors read and approved the final manuscript.

## Availability of data and materials

Raw RNA sequencing data generated in this study have been deposited in the European Nucleotide Archive (ENA) under project accession PRJEB123036. The dataset comprises 12 biological samples representing undifferentiated sheep embryonic stem cells, definitive endoderm, hepatoblast and hepatocyte-like cell differentiation stages.

## Competing interests

DR and JM are employees of Roslin Technologies Ltd. DCH is a director and co-founder of Stemnovate Limited and Stimulver ApS. These affiliations did not influence the study design, data collection, analysis, interpretation, or the decision to publish the results.

## Funding

This work was funded by Roslin Technologies and Innovate UK’s KTP programme (13974).

## Authors’ contributions

FK designed and optimized the differentiation protocol, performed the experiments, analyzed the data, prepared the figures. AK provided technical support and expertise during the initial establishment of the differentiation system. DR oversaw the project and provided scientific input. PJM and DCH conceived the project, contributed to the experimental design, data interpretation and project supervision. FK, DCH and PJM drafted the manuscript.

## Acknowledgements

The authors thank Roslin Technologies staff for discussion and technical assistance. We are also grateful to Wei-Yu Lu and Fiona Pilgrim from the University of Edinburgh and the team from the East of Scotland KTP Centre for their assistance.

