## Supplementary Data for "Generation of hepatocyte-like cells from sheep embryonic stem cells"

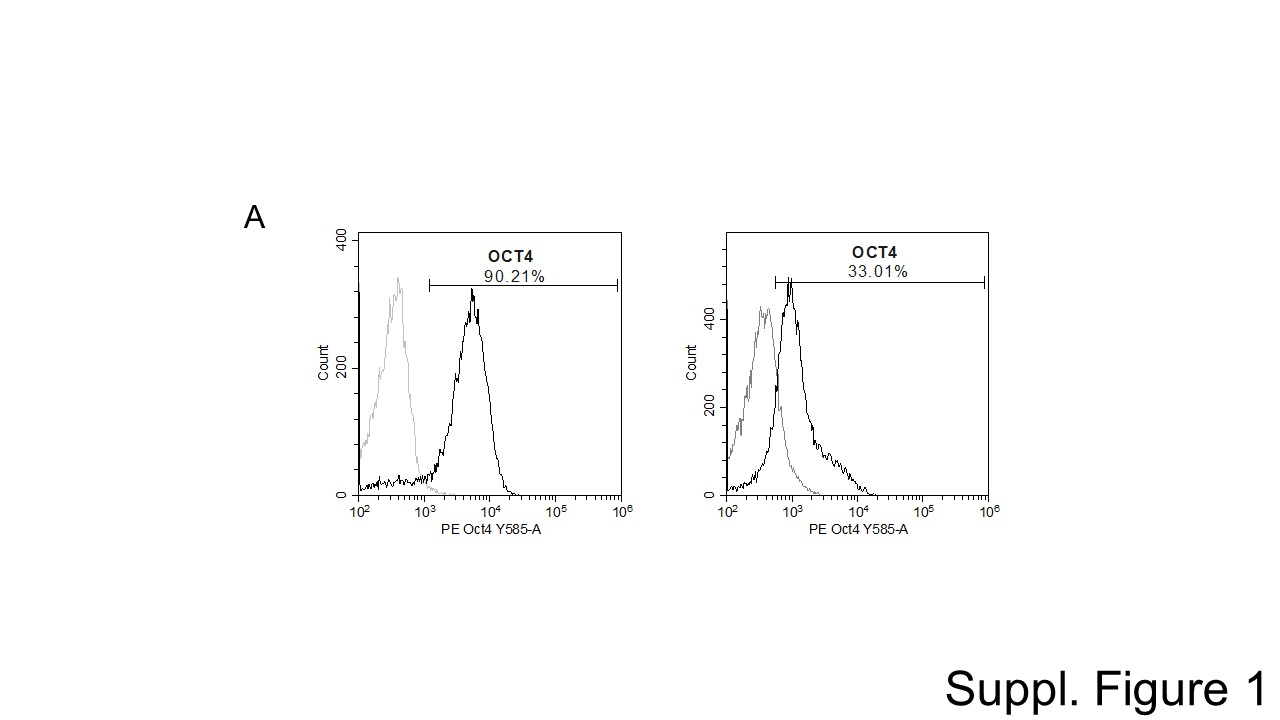

**Suppl Fig1:** Flow cytometry analysis of demonstrates a reduction in Oct4 expression in during ESC differentiation. Undifferentiated ESCs (left) and DE (right). Gray histogram represents isotype control, black histogram represents OCT4 staining.

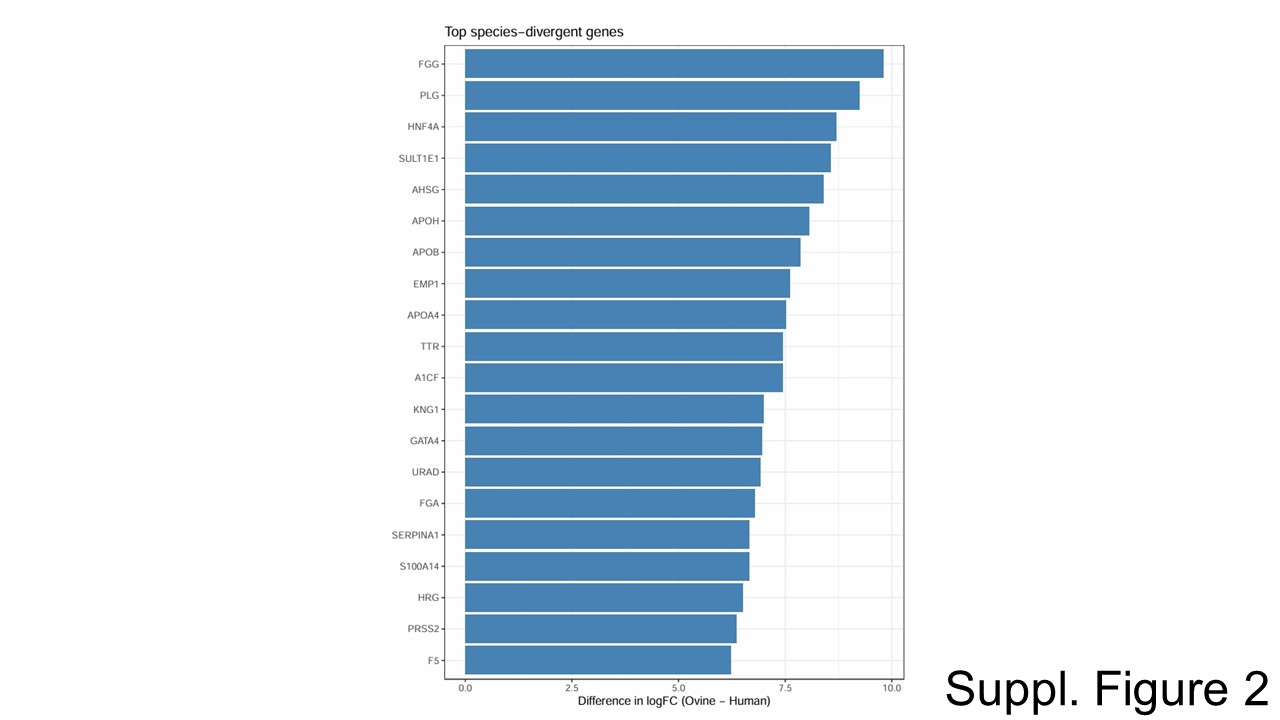

**Suppl Fig 2:** Differential gene expression summary of top 20 genes.

Top 20 differentially expressed genes (DEGs) between ovine and human HLCs.
